# Deciphering signaling specificity with interpretable deep neural networks

**DOI:** 10.1101/288647

**Authors:** Yunan Luo, Jianzhu Ma, Yang Liu, Qing Ye, Trey Ideker, Jian Peng

**Author notes:** These authors contributed equally to this manuscript (Yunan Luo, Jianzhu Ma).

## Abstract

Protein kinase phosphorylation is a prevalent post-translational modification (PTM) regulating protein function and transmitting signals throughout the cell. Defective signal transductions, which are associated with protein phosphorylation, have been revealed to link to many human diseases, such as cancer. Defining the organization of the phosphorylation-based signaling network and, in particular, identifying kinase-specific substrates can help reveal the molecular mechanism of the signaling network. Here, we present DeepSignal, a deep learning framework for predicting the substrate specificity for kinase/SH2 sequences with or without mutations. Empowered by the memory and selection mechanism of recurrent neural network, DeepSignal can identify important specificity-defining residues to predict kinase specificity and changes upon mutations. Evaluated on several public benchmark datasets, DeepSignal significantly outperforms current methods on predicting substrate specificity on both kinase and SH2 domains. Further analysis in The Cancer Genome Atlas (TCGA) demonstrated that DeepSignal is able to aggregate mutations on both kinase/SH2 domains and substrates to quantify binding specificity changes, predict cancer genes related to signaling transduction, and identify novel perturbed pathways.

**Availability:** Implementation of DeepSignal is at https://github.com/luoyunan/DeepSignal

## Introduction

Signaling networks transmit external environmental signals and mediate complex cellular communications to drive coordinated expression responses. Perturbations of this cellular signaling system have been found to underlie many common human diseases, such as diabetes, heart disease and cancers (Lahiry et al., 2010). It is critical to uncover the organization and track the dynamics of signaling networks for the purpose of therapy development.

The signaling transduction of the network is mainly performed by the protein phosphorylation, a post-translational modification (PTM) of serine, threonine or tyrosine residues, regulated by protein kinase and SH2/3 domains. The human genome encodes more than 500 unique kinases which can be organized in a hierarchy of 10 groups, 134 families and 201 subfamilies. At the molecular level, kinases regulate many cellular signaling processes by adding phosphoryl group to substrate proteins. Therefore, a critical step to understand the signaling system is to predict kinase-specific phosphorylation sites given a kinase sequence and a substrate. Despite tens of thousands phosphosites have been experimentally verified using the high-throughput mass-spectrometric techniques, our knowledge of the phosphorylation-driven signaling network is still preliminary: the upstream kinases responsible for the phosphorylation are unknown and a significant number of phosphosites are unidentified.

Many computational methods have been developed to predict phosphosites for human kinases, including Scansite (Obenauer, 2003), PredPhospho (Kim et al., 2004), NetPhosK (Blom et al., 2004), KinasePhos (Wong et al., 2007), GPS (Xue et al., 2008), Predikin (Ellis and Kobe, 2011), PSEA (Suo et al., 2014), PhosPhoPICK (Patrick et al., 2016) and PhosphoPredict (Song et al., 2017). Existing methods suffer from various limitations. First, most of these methods are kinase-specific models and share the “one-kinase-one-model” paradigm. For a given kinase, an individual model was built by utilizing the known substrate sequences of this kinase. These models are limited to those well-characterized kinases and conceptually cannot be generalized to new kinases (i.e., kinases not in training data) or kinases with mutations. Second, majority of these methods have narrow coverage and coarse resolution. For example, PhosphoPredict (Song et al., 2017) and (Li et al., 2010) can predict phosphosites for only 12 and 8 kinase families, respectively. They were trained at the kinase family level and cannot make accurate predictions for individual kinases or mutated kinases. Third, existing methods have ignored the information provided by kinase sequence, and therefore, fail to predict how the specificity would change if missense substitutions occur in the kinase sequences. Furthermore, it has been hypothesized that the substrate specificity of a kinase is not only determined by the entire domain. Instead, only a subset of residues within the domain, which are called determinants of specificity (DoS), contribute to define its specificity (Creixell et al., 2015). Several structural studies (see ref. in (Creixell et al., 2015)) have been conducted to identify the residues that are close to the binding site as potential DoS in the 3D space, since they are more likely to cooperate together to influence the binding affinity. The residues far from the binding region are usually ignored in these structure-based studies, even though they may greatly contribute to the binding specificity through long-range dependencies. An evolutionary algorithm, KINspect, was proposed in (Creixell et al., 2015) to identify the combinations of residues that can exclusively predict the specificity profiles of a kinase as potential DoS. Nevertheless, KINspect only explored a limited number (~100) of possible combinations of residues, which covers a small portion of all possible combinations. In addition, the DoS set identified by KINspect contains around only 80 residues, i.e. about 25% of a kinase domain, making the interpretation and further experimental design difficult.

To address these issues, we present DeepSignal, a deep learning based method to predict the substrate specificity of kinase domains based on the protein sequences. Unlike most of the previous methods that only focus on substrate sequences, DeepSignal takes into account the information in both kinase domain sequences and substrate peptides, and translates a kinase sequence into a binding profile (i.e. position-specific scoring matrix (PSSM)). DeepSignal employs the Long Short Term Memory (LSTM) network, a recurrent neural network with memory units, to process the kinase sequences with various lengths using a single model, enabling the learning of universal knowledge across multiple kinase domains. This model is able to identify important residues in kinase domain sequences that best explains the substrate specificity of this kinase by exploiting both long- and short-range dependencies of residues spanning over the entire kinase domain. DeepSignal can transfer the knowledge it has learned from the available kinase-substrate data in the training data to new kinases or mutated kinases, thus greatly enabling many possible analyses, which are currently limited or infeasible by existing methods.

## Results

### Overview of DeepSignal

As shown in Figure 1, the DeepSignal framework has two components, an “encoder” network and a “decoder” network, to translate a kinase sequence to its binding substrate sequence profile. In particular, the encoder is a Recurrent Neural Network (RNN) and takes a protein kinase sequence as input, scans the sequence, and extracts important combinatorial patterns of amino acids. We use the long-short term short memory (LSTM) as the basic unit of RNN. Intuitively, the LSTM unit learns when to ‘forget’ and when to ‘remember’ the long term memory. This nice property is especially helpful when both short and long range dependencies are mixed within the sequences. The deterministic information from sequences is thus encoded as an output vector by the encoder. The decoder, in conjunction with the encoder, takes this vector of hidden state as input, and outputs a sequence profile which is a distribution of 20 amino acids for each position in a substrate peptide. The model is trained to optimize the similarity between the predicted sequence profiles and those constructed from experimental data. A more formal and detailed description is provided in **Method details**.

**Figure 1.**
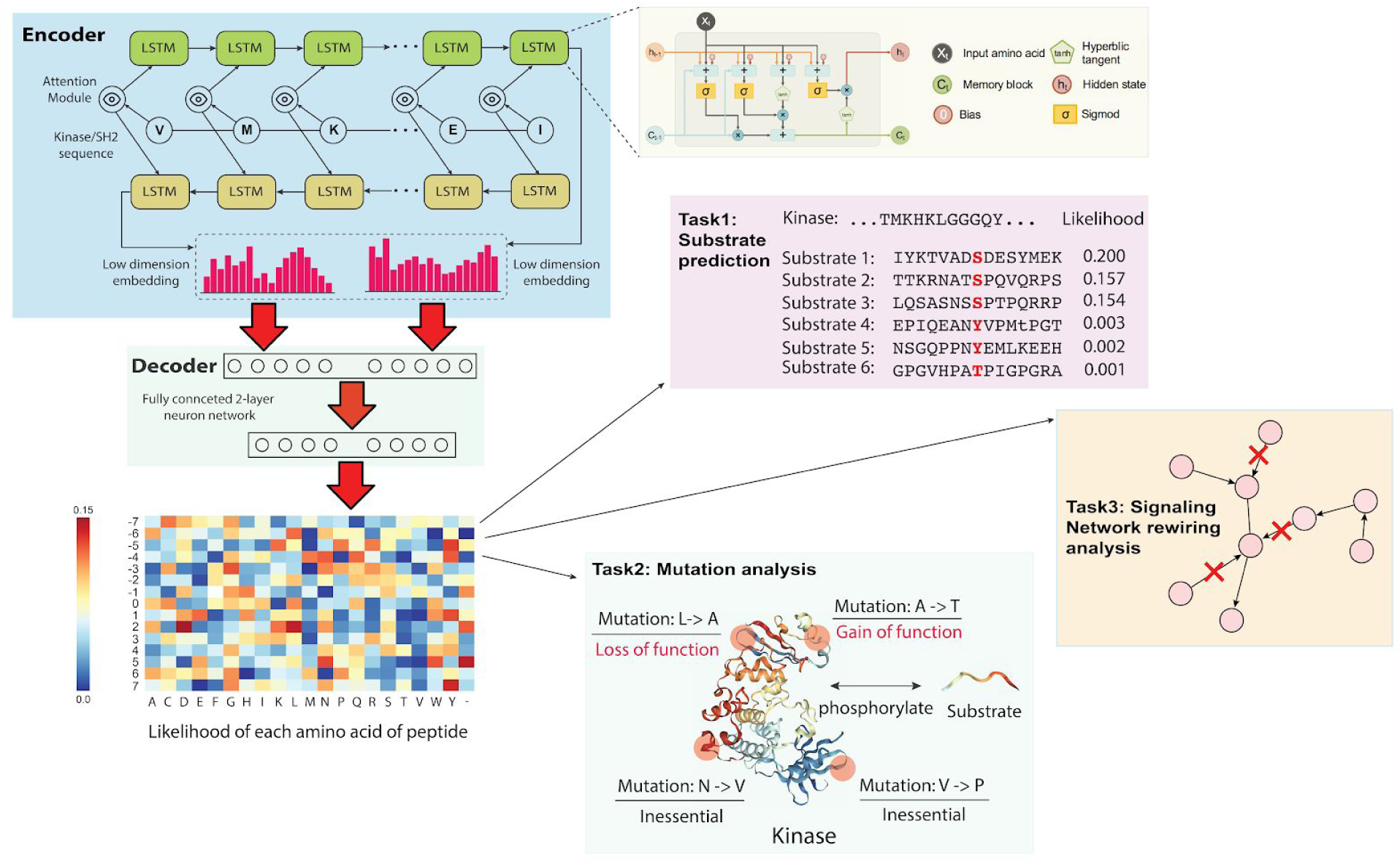
Schematic illustration of DeepSignal. DeepSignal consists of two modules, an encoder and a decoder network. The encoder network employs the bidirectional Long Short Term Memory (LSTM) network to process the input kinase (or SH2) sequence. A separate attention model is trained together to determine the feature importance. The decoder network then predicts the substrate specificity profile based on the output of attention model extracted by the encoder from the kinase sequence. A zoom-in view of the LSTM unit was also provided. Here we provide three applications of DeepSignal: 1) substrate prediction. 2) mutational analysis. 3) signaling network rewiring analysis.

### DeepSignal finds kinase specificity-determining residues

By leveraging the effective memory mechanism of LSTM, DeepSignal can automatically decide whether each residue should be enhanced or silenced based on the information extracted from protein sequence. This mechanism enables DeepSignal to efficiently explore a very large number of amino acids combinations within the kinase domains with both long- and short-range dependencies which defines the specificities between kinases and peptides. To illustrate this, we followed the setting of previous work (Creixell et al., 2015) to investigate the relationship between the 516 kinase sequence similarity and 10,181 substrate specificity similarity from several public databases (**Method details**). First, we computed the pairwise BLOSUM62 score (Henikoff and Henikoff, 1992) between two kinase domain sequences, and the pairwise negative Frobenius distance as the pairwise similarity for kinase substrate specificity. Across all the kinase-peptide binding samples, we calculated the Spearman’s correlation between the kinase sequence similarity and binding specificity similarity. We observed there is a relatively low correlation (<0.5) between these two types of similarity (Figure 2A), which suggests that not all the amino acids contribute equally to determine the substrate specificity. Next, we computed the sequence similarity on those residues identified in previous structure-based studies (Structure DoS in Figure 2A) and those predicted by KINspect, a genetic algorithm that enumerates subsets of residues whose similarity correlates to the specificity similarity (Creixell et al., 2015). As demonstrated in Figure 2A, these residues have higher correlations with the specificity, indicating that instead of the global sequence, a subset of residues potentially determines the kinase-substrate specificity. In addition, we checked whether DeepSignal is able to extract sufficient information that is encoded within the kinase sequence, by encoding the necessary dependencies among residues that determines the binding specificity. The similarity between two kinase sequences is calculated by their negative *L*_2_ distances of encoder’s outputs. DeepSignal achieved the highest correlation to the binding specificity in comparison to other methods (Figure 2A, B). This result indicates that DeepSignal can better capture the complex features and patterns in the domain sequences that determine the substrate specificity.

**Figure 2.**
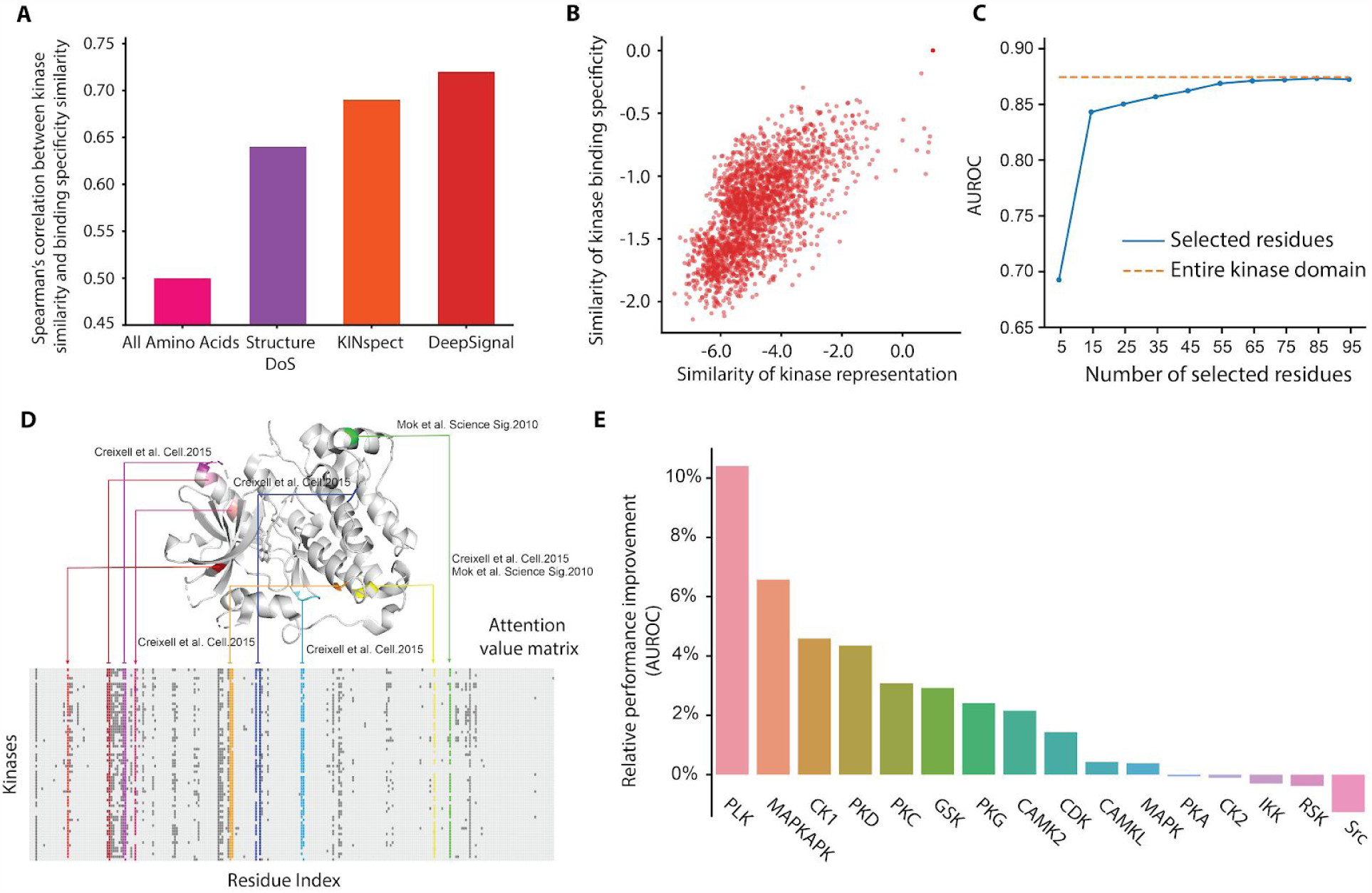
Pairwise relationships between the kinase representation similarity and kinase substrate specificity similarity. DeepSignal is able to learn a vector representation for a kinase domain that encodes the complex features determining its substrate specificity. **(A):** DeepSignal achieves higher Spearman’s correlation than the pairwise kinase domain similarity computed based on the whole domain sequences, determinants of specificity identified in previous structural studies and the KINspect algorithm. **(B):** The pairwise kinase domain similarity based on the representations learned by DeepSignal are highly correlated with the pairwise kinase substrate specificity similarity. **(C):** The prediction performance (AUROC score) of DeepSignal converges with only a small number of residues selected by the attention module. **(D):** Up: Top 15 residues selected by the attention module mapped on a reference kinase structure (PDB ID:3poz). Below: The attention value matrix learned from training data for each residue on the kinase sequence. **(E):** Relative performance improvement across 16 kinase families by including other kinase families.

Next, we studied whether the better sequence representation by DeepSignal leads to a better set of specificity-determining residues, which requires us first to exact the sequence representations. Recent progress had been made to improve the interpretability of deep learning model (Bahdanau et al., 2014; Lei et al., 2016; Ma et al., 2018). In this work, with the help of the attention model, we can extract the residue patterns from what DeepSignal learned. In particular, the attention component helps us to identify an influential set of residues to best explain the prediction. The “attention” for each residue was generated by a local neural network which considers its flanking regions on the sequence to assign an importance weight to this residue. With this attention model, we identified a subset of 15 important residues which are still able to achieve a very good prediction accuracy (with an AUC score ~0.85, comparable to the performance 0.87 trained using the full sequence) (Figure 2C, D). Among these 15 amino acids, two of them (Mok et al., 2010) had been confirmed by previous work to be important for determining binding specificity. Five residues had been annotated by KINspect (Creixell et al., 2015) as the Determinants of Specificity (DoS), and four residues are structural neighbors of them. The rest four residues are novel and might suggest new mechanisms for kinase binding. Another key observation is that, as shown in Figure 2D, important residues are almost overly distributed on the entire kinase sequence, which suggests that the determinants of the kinase binding may not be simply explained by local sequence motifs. This also explains why DeepSignal outperforms existing methods, since the RNN/LSTM model is especially suitable to mine long-range patterns.

To test the generalizability of the identified specificity-determining residues, we evaluate whether our model can transfer information across different kinase families. To explicitly demonstrate it, we restricted the testing data within only one kinase family and compared the performance of using or not using the kinases outside the family. For each kinase family with >100 peptides, as Figure 2E shows the relative performance improvement when other kinase families were included in the training data. By introducing other kinase families, DeepSignal can improve the performance on 75% (12/16) of kinase families. The AUROC scores were significantly improved on 25% of 4/16 families. Given the limited number of kinase-peptide binding pairs within each family, the ability of transferring knowledge across families alleviates overfitting and generates more robust and accurate predictions.

### Predicting kinase-specific binding sites

Armed with a better sequence representation, we compared DeepSignal (encoder + decoder) with four popular methods, GPS (Xue et al., 2008), NetPhorest (Miller et al., 2008), NetPhos (Blom et al., 2004) and MusiteDeep (Wang et al., 2017), on predicting the kinase-specific phosphosites. Since current databases contains only experimentally verified kinase-substrate pairs (positive samples), we adopted the following procedure to generate negative samples: for an experimentally verified kinase-substrate < *x*, *y* > and another randomly sampled kinase *x*′ that is not classified to the same kinase family as *x*, we added < *x*′, *y* > as a negative sample into the training and validation data. The data was split into five partitions with equal size and each partition was iteratively held out as the test set. The area under the receiver operating characteristic curve (AUROC) and the area under the precision-recall curve (AUPRC) were used as the evaluation metric to measure the performance. As shown in Figure 3A, we observed that GPS and NetPhorest have similar performance (AUPRC=~0.21), NetPhos and MusiteDeep achieved relatively poor/higher performance (AUPRC=~0.18). DeepSignal significantly outperformed these four methods by about 20% (AUPRC = 0.25). In Figure 3B, we showed that DeepSignal achieved higher AUROC scores than the other four methods (DeepSignal=0.875, GPS=0.836, NetPhorest=0.831, NetPhos=0.840 and MusiteDeep=0.818). For detail comparison, we also validated the performance of DeepSignal separately on serine and threonine (S/T) specific kinases and tyrosine (Y) specific kinases. Figure 3B showed that DeepSignal generated more accurate predictions than the other four methods on both types of kinases. These results demonstrated the ability of DeepSignal of predicting phosphosites of kinases.

**Figure 3.**
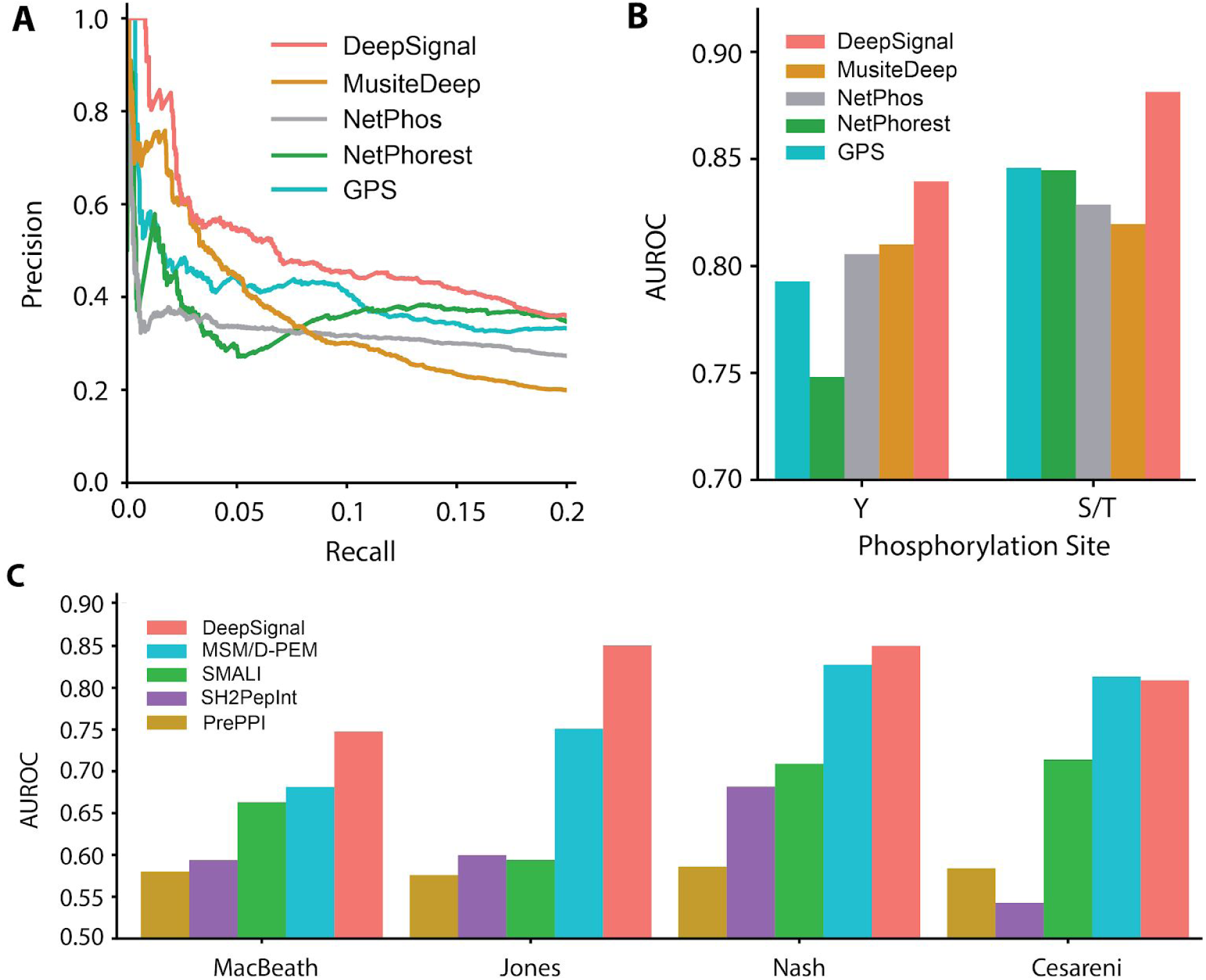
Evaluation of prediction performance on the prediction of kinase-specific phosphosites. **(A):** Precision-recall curves of five-fold cross-validation comparison of different methods. **(B):** Stratification of predicting tyrosine-specific kinases, and serine/threonine kinases, respectively. Performances were evaluated with a five-fold cross-validation and AUROC was used as the evaluation metric. **(C):** Performance of DeepSignal evaluated on four high-throughput datasets of SH2-peptide binding data and compared to four other baseline methods. Five-fold cross-validation and AUROC score were used to evaluate each method. Note that the released model of MSM/D-PEM has been trained on all the data of each dataset, while our method was trained on the training data only in the cross-validation test.

### Application to SH2 domains

In addition to the kinase domains, we also applied DeepSignal to predict the substrate specificity on SH2 domain proteins, which also play an important regulatory role in cellular signaling. Here, we collected the binding datasets from four high-throughput studies: MacBeath (Koytiger et al., 2013), Jones (Hause et al., 2012), Nash (Liu et al., 2010), and Cesareni (Tinti et al., 2013). We compared DeepSignal with other three state-of-the-art SH2-peptide binding prediction methods: SMALI (Li et al., 2008), SH2PepInt (Kundu et al., 2013) and MSM/D-PEM (AlQuraishi et al., 2014), and one general protein-protein interaction prediction method, PrePPI (Zhang et al., 2013).

We observed that DeepSignal outperformed the baseline methods in 93.8% (15/16) of the head-to-head comparisons (Figure 3C). The improvement achieved by DeepSignal was substantial. For example, on the Jones datasets, the AUROC score of DeepSignal is ~0.85, while all the other methods have AUROC score < 0.75. The only one exception is the comparison between our method and MSM/D-PEM on the Cesareni dataset, on which the two methods have comparable performance (DeepSignal=0.809 and MSM/D-PEM=0.813). However, it should be noted that although both methods were tested on the same testing dataset held out in the five-fold cross-validation, the released MSM/D-PEM model was trained with all the binding data on each dataset, while our method was trained only on 80% of the training data every time in cross-validation. Therefore, MSM/D-PEM has the advantage of the access to test data and achieved the comparable performance to ours. Overall, these experimental results suggested that the application of DeepSignal to other protein binding prediction problems, such the peptide specificity of SH2 domains.

### Identification of cancer mutations that dis-regulate signaling networks

To study the impact of mutations on cancer, we applied DeepSignal to construct the signaling network using only the protein primary sequences of 16,254 proteins, including 307 kinase domains, 112 SH2 domains and 190,427 phosphoproteins across 18 cancer types (**Method details**). For each cancer type, we mapped all the missense mutations from TCGA on the protein sequences, including 6,286 mutations on kinase domains, 776 mutations on SH2 domains and 37,996 mutations on phosphoproteins. For each node (gene) and edge (gene interactions) in the signaling network, we define two scores *P*_*perturb–node*_ and *P*_*perturb–edge*_ to quantify the likelihood of whether the binding activity being activated or disrupted (**Method details**). As shown in Figure 4A, B, the tumor networks exhibited strong enrichment for known cancer genes (Iorio et al., 2016). In particular, we observed the edge-based perturbation score *P*_*perturb–edge*_ is much more sensitive in terms of ranking cancer genes compared to node-based perturbation *P*_*perturb–node*_. The reason is that *P*_*perturb–edge*_ captures each individual rewiring of signaling transduction network while *P*_*perturb–node*_ can only quantify the marginal effect over all the possible network rewiring. Figure 4A shows that for the top 50 edges ranked by *P*_*perturb–edge*_. DeepSignal can significantly improve the efficiency of detecting cancer related genes in comparison to MSM/D-PEM for majority of cancer types. DeepSignal achieves superior performance on liver cancer (LIHC), uterine cancer (UCEC), bladder cancer (BLCA) and colon and rectal cancer (COADREAD). For instance, for LIHC, DeepSignal was able to detect 55% of the known cancer genes by considering the mutation impacts on signaling pathway. To further study these 4 cancer types, we calculated the efficiency of detecting cancer genes by choosing different ranking thresholds. As shown in Figure 4C-J the known cancer genes are constantly ranked higher by DeepSignal when compared to MSM/D-PEM, suggesting that our method is able to quantitatively measure the signaling perturbations that are relevant to important cancer signaling pathways.

**Figure 4.**
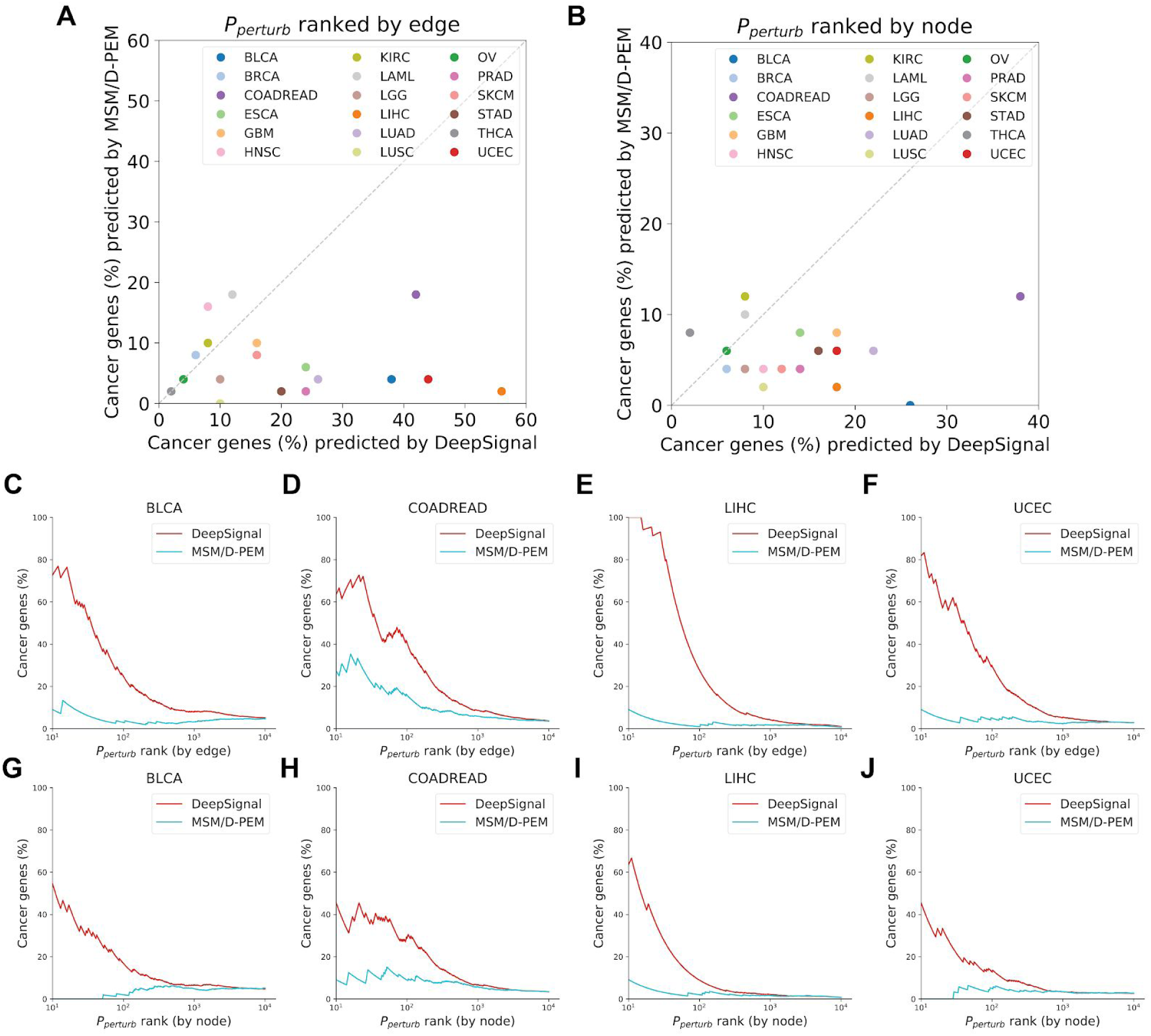
Comparison of known cancer gene detection between DeepSignal and MSM/D-PEM. X-axis and Y-axis are the percentage of genes known to be involved in a particular cancer type within the top 50 ranked proteins by DeepSignal and MSM/D-PEM, respectively, according to the predicted specificity changes. Rankings were performed on the basis of **(A)** edges and **(B)** nodes. The comparison of top genes identified by DeepSignal with known cancer genes is also provided. **(C)-(F):** The enrichment analysis of top genes ranked by the edge-based perturbation score. **(G)-(J):** The enrichment analysis of top genes ranked by the node-based perturbation score.

We further chose three of these significantly perturbed signaling pathways to demonstrate how DeepSignal could be applied to facilitate biologists and clinicians to study the mechanism of cancer. In Figure S1A, we focus on the perturbed edges associated with gene *CTNNB1* in *UCEC*. *CTNNB1* is ranked 1st and 2nd by the *P*_*perturb–edge*_ and *P*_*perturb–node*_ score, respectively. *CTNNB1* is known as a very important component of adhesion junctions which are necessary for the creation and maintenance of epithelial cell layers by regulating cell growth and adhesion between cells. Computational studies showed that *CTNNB1* mutations correlated with the upregulation and downregulation of multiple well-known cancer genes (Ding et al., 2015) but the mechanism is unclear. In Figure S1B, we focused on the perturbed edges associated with gene *PTEN* in glioblastoma (GBM), which is ranked 3rd and 6th by *P*_*perturb–edge*_ and *P*_*perturb–node*_ score, respectively. *PTEN* was identified as a tumor suppressor that is mutated in a large number of tumors at high frequency by negatively regulating AKT/PKB signaling pathway (Yang et al., 2017). Figure S1B shows its interactions with other very important cancer gene *FGFR2* and *FGFR3* are disrupted, suggesting the functional role of *PTEN* involved in cancer. We also checked the perturbed edges associated with gene *SMAD4* in *LUAD*, which is ranked 4th and 10th by *P*_*perturb–edge*_ and *P*_*perturb–node*_ score, respectively. *SMAD4* plays a central role in the balance between atrophy and hypertrophy, which is not only associated with cancer but also associated other disease such as de Myhre Syndrome and Polyposis, Juvenile Intestinal (Haeger et al., 2016). As shown in Figure S1C, DeepSignal predicts that the mutations of *SMAD4* will only lead to three significant network rewiring events which dramatically reduce the space for pharmacist to search for new drug targets to cure lung cancer.

## Conclusion

We introduced DeepSignal, a deep learning based method for predicting the substrate specificity of kinase/SH2 domains. DeepSignal consists of two components, an encoder network that encodes the sequence features to a compact expressive low dimensional vector that can better explain the substrate specificity, and a decoder network that translates the vector into a specificity profile (e.g., PSSM). With the powerful memory mechanism of the LSTM neural network architecture, we are capable of handling variable-length kinase domain sequences and effectively translate the protein sequence of a kinase into a substrate specificity profile. Experiments demonstrated that DeepSignal could achieved improved performance than existing methods. Further interpretation of DeepSignal model showed its strong predicting power came from better capturing different complicated and long-range amino acid patterns. Moreover, by analyzing phosphorylation-related missense coding mutations in TCGA data, DeepSignal can propose new candidate pathways that potentially disrupt the normal signaling processes of cell caused by cancer mutations, which may eventually help study new treatment methods to cancer patients.

DeepSignal is a versatile method and is applicable to various analyses related to kinase substrate specificity or signaling. For example, predicting the binding likelihood of a given kinase and substrate, quantifying the change of substrate specificity caused by mutations in the kinase sequence, and predicting the impact of mutations in peptides on phosphorylation events (e.g., phosphosites disruption, creation or re-wiring). Given the transferability of its predictive power, DeepSignal can be applied to predict new substrate sequences for a kinase that have few known substrates or with mutations, which will reveal important functional disruptions caused by mutations in human diseases.

## STAR Methods

### Contact for Reagent and Resource Sharing

Jian Peng, Department of Computer Science, University of Illinois at Urbana-Champaign, Urbana, IL, USA.

## Method Details

### Dataset preparation

#### Kinase and SH2 domain sequences

Protein sequences of human kinome were downloaded from the http://kinase.com repository, which provides the domain sequence and additional functional information for 516 known human kinases. These domain sequences were searched using HHblits (Remmert et al., 2011) and re-aligned using HHalign (Söding et al., 2005) with their default parameter settings. We also collected 112 SH2 domain sequences whose alignments are from a previous work (AlQuraishi et al., 2014), in which the SH2 domains were aligned with SH2-peptide structural complexes.

#### Substrate peptides

Phosphorylation sites data were extracted from the PhosphoSitePlus database (Hornbeck et al., 2012), which contains 17,272 experimentally validated phosphosites. Each phosphosite is represented by a flanking sequence of length 15 centered around the phosphosite. We extracted the human phosphosites and removed duplicates entries, leaving 10,181 non-redundant human phosphosites, spanning 307 human kinase domains. For SH2 domain, we obtained the collected dataset used in (AlQuraishi et al., 2014), which is the union of four high-throughput binding data (“MacBeath” (Koytiger et al., 2013), “Jones” (Hause et al., 2012), “Nash” (Liu et al., 2010), and “Cesareni” (Tinti et al., 2013)) of SH2 domain-phosphosite binding, including 5,016 phosphotyrosine peptide sequences spanning 111 SH2 domains.

#### TCGA dataset

Somatic missense single-nucleotide variants of 24 cancer types and 4,954 patients, including 5,196 samples and 19,934 genes, from The Cancer Genome Atlas (TCGA) were downloaded from the Broad GDAC Firehose Hub (http://gdac.broadinstitute.org, February 11th 2016).

### The DeepSignal Framework

Our method applies the recurrent neural network (RNN) (Hochreiter and Schmidhuber, 1997), which is a deep learning architecture that has demonstrated its applicability in various sequence-based tasks, e.g., natural language translation (Wu et al., 2016), speech recognition (Graves et al., 2013)] and computational biology (Min et al., 2017). RNNs are able to take a protein sequence with an arbitrary length as input and maintain a hidden/memory state. Given a protein kinase sequence *x* with length *n*, at each position *t*, the RNN receives the *t*-th amino acid *x*_*t*_ of the kinase sequence, and updates the hidden state using the following transition function, which is the composition of an affine transformation followed by a non-linear function.

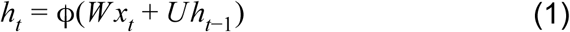

where *h*_*t*_ is the hidden state at position *t*, ϕ(⋅) is a non-linear function (e.g., *tanh*(⋅)), and *W* and *U* are the parameters of the RNN. In this work, we implemented the RNN using the Long Short Term Memory (LSTM) units. LSTM is an improved version of the RNN defined in Equation 1, which can better capture long-range dependencies by choosing proper information to forget, and at the same time benefit mathematical optimization. Intuitively, the hidden state *h*_*t*_ encodes the “internal memory” of the RNN up to position *t* and summarizes the long-short range dependencies information over multiple distance scales. In this work, we use a bidirectional LSTM (BiLSTM) (Schuster and Paliwal, 1997) that consists of two LSTMs, with one scanning the sequence forward and the other scanning backward. For simplicity, we use *h*_*t*_ to represent the concatenation of the hidden states of the forward and backward LSTMs at position *t*, i.e.,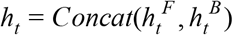, where 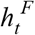 and 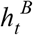 are the hidden states of the forward and backward LSTMs, respectively. To identify the residues in the kinase sequence that contribute the most to the determination of specificity (the so-called determinants of specificity, or DoS), we employed the attention mechanism to our DeepSignal model, which assigns an attention weight to each residue in the kinase sequence. The weights provided by the attention model were learned during the training process such that residues which are more critical in predicting the substrate specificity will receive higher weights. The attention weight of the *t*-th amino acid *x*_*t*_ was calculated as follows,

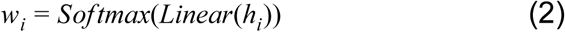

We then used the linear combination of all hidden states *h*_1_, *h*_2_, …, *h*_*n*_, weighted by the attention weights, as a context vector *c*_*i*_ to explicitly capture the information provided by each residue,

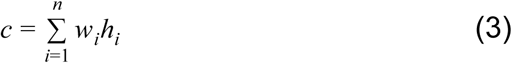

Next, we concatenated the context vector and the final hidden state *h*_*n*_ to obtain an attention vector *v* as the output of the encoder,

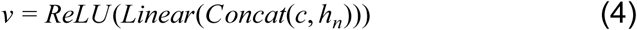

Given the kinase representation generated by the encoder, the decoder aims to predict the substrate specificity for this kinase. Specifically, for each position in the substrate peptide, the decoder will learn to predict a probability that an amino acid *a*∈Σ will occurs at that position, where Σ is the set of all the 20 possible amino acids. For this purpose, we use multiple independent two-layer neural network. That is, there is a different neural network for each position in the peptide, to learn the specificity at each position. In particular, for the *k*-th position in the peptide, the two-layer neural network is defined as,

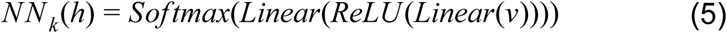

where *h*_*n*_ is the output of the encoder, *Linear*(⋅) is an affine transformation as *Linear*(*x*) = *Wx* + *b*; *ReLU* (⋅) is the Rectified Linear activation Unit (Nair and Hinton, 2010) used to imitate the neuron activation and defined as *ReLU* (*x*) = *max*(*x*, 0); the softmax function defined as 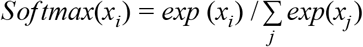 is used to normalize the output to be a valid probability distribution. The output of *NN_k_* (*h*_*n*_), denoted as *p*_*k*_, is a 20-dimension vector indexed by 20 amino acids, in which the entry *p*_*k*_ (*a*) is the probability that the amino acid *a* occurs at the *k*-th position in the peptide. Given a set of training examples {(*x*^(*i*)^, *y*^(*i*)^)}, where _*x*_(*i*) is a kinase sequence and _*y*_(*i*) is a peptide sequence, centering on the phosphorylation site, the loss function of our model is defined as the sum of negative log-likelihood (*NLL*), i.e.,

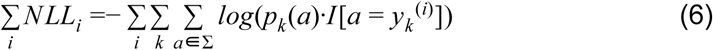

We used the stochastic gradient descent (error backpropagation + ADAM (Kingma and Ba, 2014)) to minimize the negative log-likelihood.

### Signaling Perturbation Score

We construct a signaling network from TCGA data, in which nodes represent kinase proteins, SH2 proteins and phosphoproteins that contains somatic missense single-nucleotide mutations, and edges represent the phosphorylation relationships between wild-type kinase/SH2 proteins and wild-type peptide. Taking the sequence *x* of the wild-type kinase/SH2 as input, DeepSignal predicts the substrate specificity as a sequence profile, i.e., a probability distribution in which *p*_*k*_ (*a*) gives the probability that amino acid *a* occurs at the *k*-th position in the peptide. This distribution can be used to evaluate the likelihood *L*_*w*_ that a wild-type peptide (with sequence *y*) will by phosphorylated by the kinase/SH2,

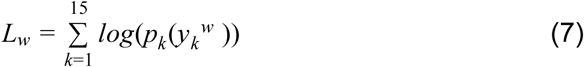

Now let *x*′ and *y*′ be the corresponding mutant sequences of the kinase/SH2 protein and the peptide (it is also possible only one of the kinase/SH2 protein and peptide was mutated). Taking *x*′ as input, DeepSignal predicts the substrate specificity *p*′_*k*_ (*a*) for the mutant kinase/SH2, and the likelihood *L*_*m*_ that a mutant peptide will by phosphorylated is given by,

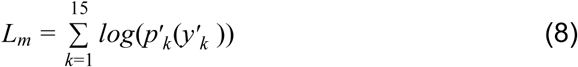

The perturbation score of the edge *e* between the kinase/SH2 *x* and the peptide *y* is then defined as,

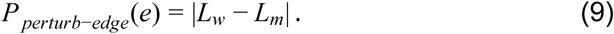

The perturbation score of a node *P*_*perturb–node*_ is defined as the aggregation of the *P*_*perturb–edge*_ scores of all edges associated with this node, i.e.,

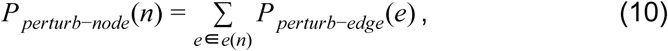

where *e*(*n*) is the set of edges that are incident to node *n*.

### Baseline methods

For kinase substrate specificity prediction, we compared DeepSignal with four state-of-the-art methods (GPS (Xue et al., 2008), NetPhorest (Miller et al., 2008), NetPhos (Blom et al., 2004) and MusiteDeep (Wang et al., 2017)). For SH2 substrate specificity prediction, we compared DeepSignal to three existing SH2 models (SMALI (Li et al., 2008), SH2PepInt (Kundu et al., 2013) and MSM/D (AlQuraishi et al., 2014)) and one general protein-protein interaction method (PrePPI (Zhang et al., 2013)). Among these methods, GPS, SMALI and SH2PepInt are kinase/SH2-specific, which means a specific model is trained specifically for a given kinase/SH2 and the model can only predict the phosphosites for that kinase/SH2. NetPhorest integrates specificity profiles or PSSM derived from Positional Scanning Peptide Library experiments and uses them to predict the phosphosites of a kinase. MSM/D uses a multiscale framework to model the SH2 specificity and derives the position energy matrices (PEMs) to describe the binding selectivity. In our experiments, the prediction method based on PEMs derived from MSM/D is denoted as MSM/D-PEM. Note that the pre-built specificity profiles of both NetPhorest and MSM/D-PEM were derived from all available data, which may overlap with part of the test data in our cross-validation process. Therefore, we gave advantages to the existing baseline methods but not DeepSignal in the cross-validation comparisons. For NetPhos, we ran their web services with default parameters. MusiteDeep provided pre-trained models for both kinase-specific phosphosites and general phosphosites prediction. For a given kinase, if its pre-trained kinase-specific model has been provided by MusiteDeep, we used this kinase-specific model to predict the phosphosites, otherwise we used the general phosphosites prediction model.

### Parameter settings and training details

The hyperparameters of DeepSignal were selected using an inner-loop 5 fold cross-validation on the training data only. Specifically, the encoder is a BiLSTM with 512 hidden units (i.e., the dimension of the hidden state vector) and the decoder is a 2-layer neural network with 512 hidden units. We trained the model for 500 epochs with batch size 512. The learning rate was set to 0.001. Several regularization techniques, e.g., dropout (Srivastava et al., 2014) and early stopping (Goodfellow et al., 2016), were used to avoid overfitting. Kinases with less than 30 known substrates or with 20 known phosphorylation sites were filtered out in the training data. The model was implemented in PyTorch and trained on NVIDIA Titan X GPUs. The training process takes 15s for one epoch and 500 epochs to converge.

## DATA AND SOFTWARE AVAILABILITY

A pytorch implementation of DeepSignal, including both training and predicting codes and a benchmark dataset are available for download at: https://github.com/luoyunan/DeepSignal.

---

**Figure S1.**
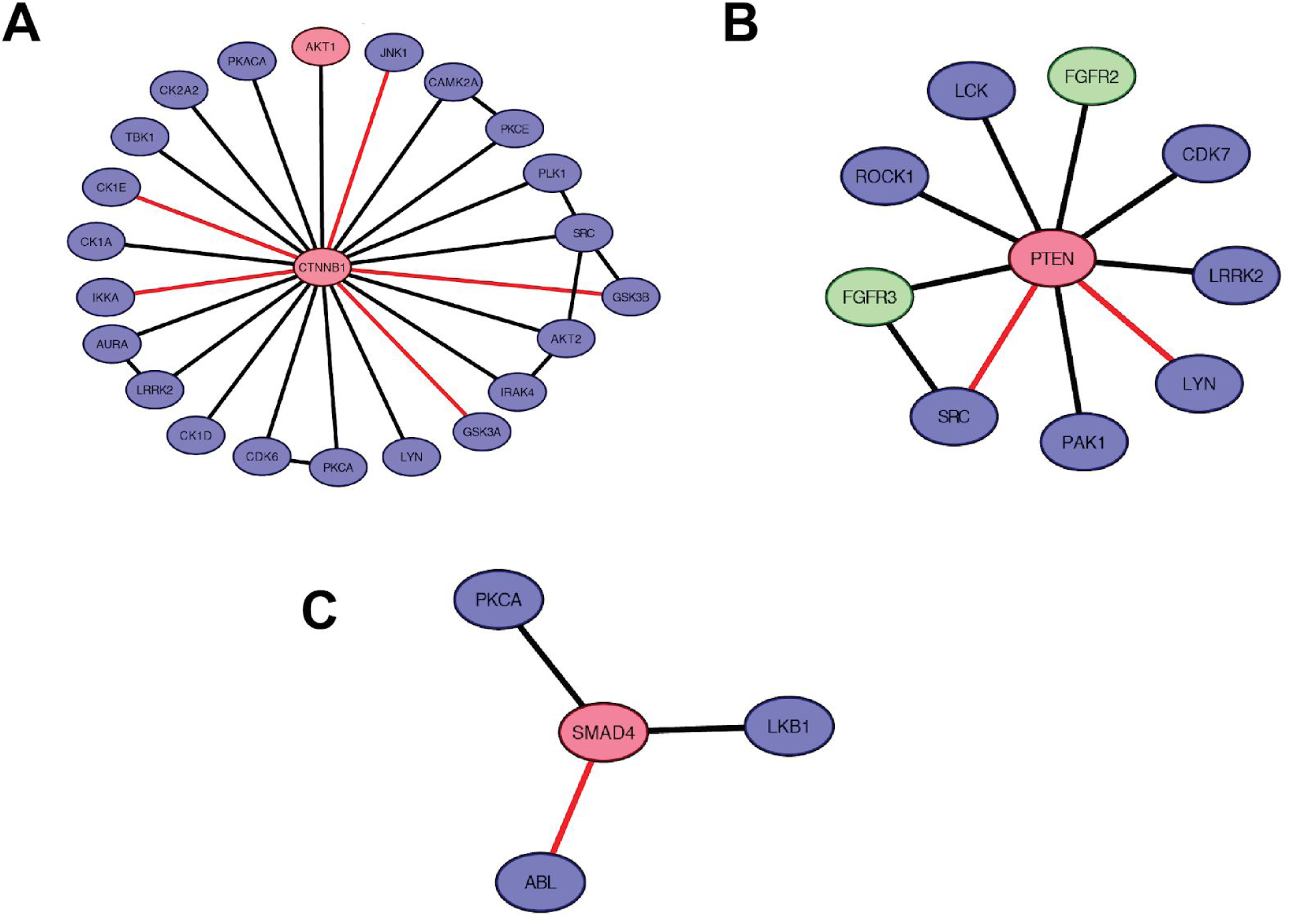
DeepSignal discovers new perturbed pathways related to cancer including **(A):** CTNNB1 pathway in UCEC, **(B):** PTEN pathway in GBM and **(C):** SMAD4 pathway in LUAD. Black edges represent the wild-type interactions between Kinase/SH2 domain and phosphoproteins. Red edges represent loss of functions predicted by DeepSignal. Red color of nodes represents known cancer genes for the corresponding cancer type and green color of nodes represents known cancer genes for other cancer types.

